# A pro-apoptotic selection strategy enables CRISPR screening in mosquitoes and identifies Lachesin as a chikungunya virus entry factor

**DOI:** 10.64898/2026.07.31.742019

**Authors:** Anja C. M. de Bruin, Enzo Mameli, Lathika Valliyott, Yanhui Hu, Raghuvir Viswanatha, Jesse S. Plung, Wanyu Li, Jonathan Abraham, Andres Merits, Nicholas Ariotti, Norbert Perrimon, Gisa Gerold

**Affiliations:** Institute of Virology, Medical University of Innsbruck, Innsbruck, Austria; Department of Genetics, Blavatnik Institute, Harvard Medical School, Boston, MA, USA; Institute for Molecular Bioscience, The University of Queensland, St Lucia, Queensland, Australia; Department of Microbiology, Blavatnik Institute, Harvard Medical School, Boston, MA, USA; Howard Hughes Medical Institute, Boston, MA, USA; Institute of Bioengineering, University of Tartu, Tartu, Estonia

## Abstract

Chikungunya virus (CHIKV) is a re-emerging mosquito-borne alphavirus that is transmitted primarily by *Aedes aegypti* and *Aedes albopictus* mosquitoes. CHIKV infection can result in debilitating arthritis-like symptoms in humans. While determinants of CHIKV host cell entry into mammalian cells are known, their equivalents in mosquito cells remain largely elusive. To identify CHIKV entry factors, we performed a membrane-focused genome-scale CRISPR loss-of-function screen in *Aedes albopictus* mosquito cells. To enable selection of refractory cells, we engineered CHIKV to express the pro-apoptotic *Drosophila* Reaper protein. CHIKV-Reaper induced robust cell death in mosquito cells with only modest viral fitness loss. Using this selection strategy together with an *Aedes albopictus* C6/36-based CRISPR screening platform, we identified glycosylphosphatidylinositol (GPI)-anchored cell surface proteins and multiple enzymes involved in the GPI-anchor biosynthesis pathway as proviral candidates. Ectopic expression of the GPI-anchored cell-adhesion protein Lachesin rendered refractory mammalian cells susceptible to CHIKV and the related arthritogenic alphaviruses Semliki Forest virus and Ross River virus. In contrast, Lachesin-expressing cells remained refractory to the encephalitic alphavirus Venezuelan equine encephalitis virus. We demonstrate that Lachesin is essential for CHIKV infection in both *Aedes aegypti* and Aedes albopictus cells, as confirmed by gene silencing. Here, we identify Lachesin as a critical candidate entry receptor for CHIKV and establish pro-apoptotic arboviruses as a powerful and versatile strategy for functional CRISPR screening in mosquito cells.

## Introduction

Alphaviruses (family *Togaviridae*) are small enveloped positive-sense RNA viruses of which most are transmitted between vertebrates by mosquitoes (1). These arthropod-borne viruses, i.e. arboviruses, can be subdivided into Old World and New World alphaviruses, with the former primarily causing arthritogenic disease and the latter encephalitic disease (2, 3). Chikungunya virus (CHIKV) is a re-emerging Old World alphavirus that causes outbreaks in tropical and sub-tropical regions and more recently in Europe, China, and North America. In 2025, the European mainland reported more than 1,100 cases of CHIKV disease, while China recorded more than 16,000 cases (4-6). In addition to causing long-lasting debilitating arthritis-like symptoms (7), CHIKV has been associated with sporadic neurological complications and death (8). CHIKV belongs to the Semliki Forest (SF) serocomplex, which, in addition to the eponymous Semliki Forest virus (SFV), includes the clinically relevant o’nyong’nyong virus (ONNV), Ross River virus (RRV), and Mayaro virus (MAYV), amongst other non-human pathogenic alphaviruses (9). The most important New World encephalitis-causing alphaviruses are Venezuelan equine encephalitis virus (VEEV), eastern equine encephalitis virus (EEEV), and western equine encephalitis virus (WEEV).

During explosive outbreaks, CHIKV circulates between anthropophilic mosquitoes and humans in an urban epidemic transmission cycle (10). Before a large outbreak in the Indian Ocean in 2004, *Aedes* (*Ae*.) *aegypti* was the major mosquito vector transmitting CHIKV (11-13). Since then, CHIKV acquired adaptive mutations in its 11.8 kb genome, and these facilitated viral amplification in the globally invasive *Ae. albopictus* mosquito (14-16). In mosquitoes, alphaviruses share a common amplification route in which virus is ingested with an infectious blood meal and infects the midgut epithelium. After amplification in the midgut, the virus crosses the midgut barrier, disseminates through the haemocoel, and eventually reaches the salivary glands, enabling transmission to the next host via saliva injection during blood feeding. Despite the importance of mosquito vectors for transmission, molecular determinants that drive CHIKV infection in mosquitoes remain largely elusive. In particular, the specific mosquito surface proteins hijacked by the virus to enter cells have not been identified.

During mammalian cell entry, alphaviruses contact target cells through engagement of general attachment factors such as glycosaminoglycans and phosphatidylserine (PS)-receptors (17-19). Internalisation occurs upon binding of the virion to a bona fide entry receptor. The viral surface glycoproteins that facilitate receptor-mediated uptake are E1 and E2, which are organised as 80 trimers of E1-E2 heterodimers per virus particle (20). Whereas E2 is surface exposed, E1 is partially obscured and harbours the hydrophobic fusion loop. Upon virion internalisation, acidification of the endosomal lumen triggers dissociation of E1 and E2, followed by conformational changes that result in exposure of the fusion loop and membrane fusion (21-23). In mammalian cells, Matrix Remodelling-Associated protein 8 (MXRA8) has been identified as entry receptor for CHIKV and related arthritogenic alphaviruses (24, 25), Very Low Density Lipoprotein Receptor (VLDLR) and Apolipoprotein E receptor 2 (ApoER2) for SFV, EEEV, Sindbis virus (SINV), and certain WEEV strains (26), Low-Density Lipoprotein Receptor Class A Domain-Containing 3 (LDLRAD3) for VEEV (27), and Protocadherin-10 (PCDH10) for WEEV (28). In *Drosophila melanogaster*, natural resistance-associated macrophage protein (NRAMP) has been described as a SINV entry factor (29).

No orthologues of MXRA8 have been identified in mosquitoes and the factors that mediate CHIKV entry into mosquito cells are yet unknown. In contrast, mosquito orthologues of the human SFV and EEEV receptor VLDLR exist (26). Genome-scale CRISPR/Cas9 screening technologies have proved instrumental in identifying human alphavirus receptors. Here, we develop a methodology that enables CRISPR/Cas9 knockout screening in mosquito cells (30, 31) together with a novel selection strategy. Specifically, we engineer a full-length infectious CHIKV to encode the pro-apoptotic *D. melanogaster* Reaper protein, thus converting infection into a survival-based selection assay, and use this virus to identify host dependency factors in *Ae. albopictus* C6/36 cells. Reaper expression upon infection enabled stringent positive selection of CHIKV-refractory cells in our loss-of-function screen. Using this approach, we identified the glycosylphosphatidylinositol (GPI)-anchored immunoglobulin superfamily (IgSF) protein Lachesin, together with multiple enzymes required for GPI-anchor biosynthesis, as key mediators of CHIKV entry into *Ae. albopictus* and *Ae. aegypti* mosquito cells.

## Results

### Design and characterisation of pro-apoptotic CHIKV

A genome-scale CRISPR screen requires the selection of cells based on a specific phenotype. In the loss-of-function screen developed here, we aimed to correlate the genetic ablation of a host dependency factor with a loss of susceptibility to CHIKV. Generally, selection of uninfected cells is accomplished by fluorescence-activated cell sorting (FACS), magnetic-activated cell sorting (MACS), or by survival-based selection (32). Alphaviruses establish persistent infection in mosquito cells without causing overt cytopathic effects (33-36), thereby precluding survival-based selection. To overcome this limitation, we generated a pro-apoptotic CHIKV. An East-Central-South African (ECSA) and Indian Ocean Lineage (IOL) CHIKV strain (OPY1 LR2006) was engineered to encode the pro-apoptotic Reaper protein from *D. melanogaster*. Reaper is a negative regulator of the Inhibitor of Apoptosis (IAP) proteins, and its overexpression therefore causes apoptosis in both insect and human cells (37). Reaper is negatively regulated through IAP-mediated ubiquitination of lysine residues. Five lysine-to-arginine substitutions render Reaper ubiquitination-resistant (RprKR), thus preventing the negative feedback loop (37). In previous studies, SINV was engineered to express RprKR to assess the effect of apoptosis on intrinsic SINV transmission in adult mosquitoes (38, 39). Here, we engineered and compared two CHIKVs (**Fig 1A**); one harbouring RprKR under a subgenomic (SG) promoter (2SG-RprKR) and one in which RprKR was embedded in the structural open reading frame (ORF), flanked by a ubiquitin cleavage site and a 2A ribosomal skipping sequence (RprKRorf) (39, 40). The latter strategy is associated with improved viral genome stability *in vivo* (39). Importantly, both viruses display E1 and E2 glycoproteins identical to WT CHIKV, enabling their use in entry factor CRISPR screening.

**Figure 1.**
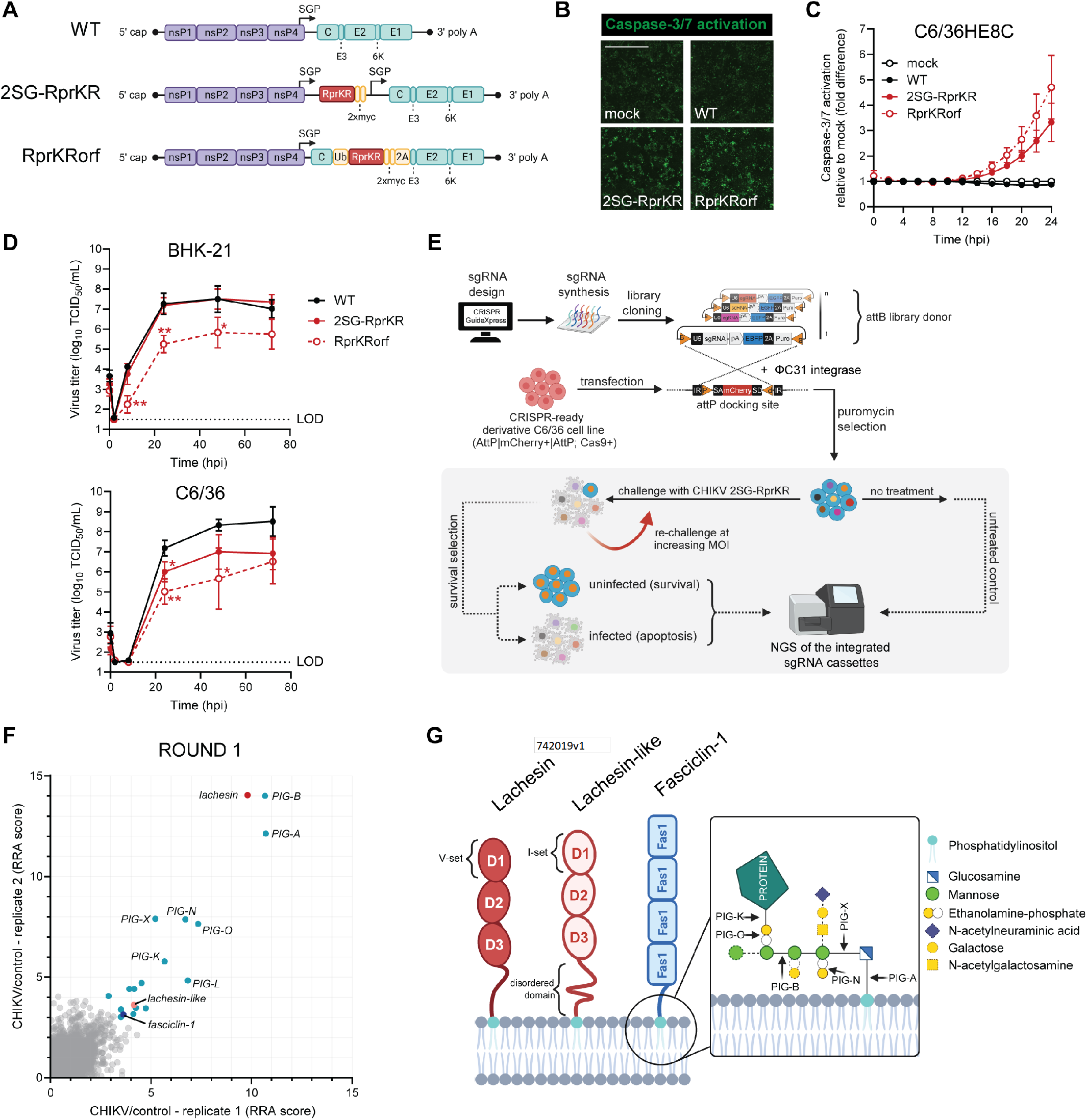
Genome-scale CRISPR screen in *Ae. albopictus* cells using a pro-apoptotic CHIKV. **(A)** Design of CHIKV (ECSA lineage - LR2006 OPY1) variants expressing *D. melanogaster* Reaper, modified with lysine-to-arginine substitutions to avoid degradation and increase apoptotic activity (RprKR). RprKR sequence is located downstream of a subgenomic promoter (SGP) in a virus designated CHIKV 2SG-RprKR, or incorporated in the structural ORF of a virus designated CHIKV RprKRorf, flanked by ubiquitin (Ub) and ribosomal skip sites (2A). WT = wild-type. **(B)** Representative images of C6/36HE8-Ub::Cas9-2A-Neo cells (C6/36HE8C) incubated with caspase-3/7 activation dye, inoculated with indicated CHIKV variants at MOI:1. Fluorescence was assessed by live cell imaging at 24 hpi. Scale bar = 400 µm. **(C)** Quantification of caspase-3/7 activation over time in C6/36HE8C cells inoculated with CHIKV at MOI:1, normalised to mock-inoculated cells. Data are mean ± SEM of three biological replicates. **(D)** Replication kinetics of CHIKV variants in mammalian BHK-21 cells or C6/36 cells. Cells were inoculated at MOI:0.01 and infectious virus titers in the supernatant were determined by endpoint titration in BHK-21 cells. LOD = limit of detection. Data are mean ± SD of three biological replicates. One-way ANOVA with unpaired t-tests compared to WT. **(E)** Schematic for pro-apoptotic CHIKV CRISPR screen. CRISPR GuideXpress was used to design a genome-scale sgRNA library that consisted of 47,677 sgRNAs targeting 6,361 genes. The library was cloned into the attB donor library vector and delivered to C6/36HE8-Ub::Cas9-2A-Neo cells via ΦC31 recombination-mediated cassette exchange to yield a pool of knockout cells. After 30 days of puromycin selection, the knockout pool was challenged with CHIKV 2SG-RprKR at MOI:0.001. At 7 dpi, surviving cells were expanded and re-challenged three times at increasing MOI. Genomic DNA from surviving cell fractions and from an uninfected knockout-pool input control was used to PCR-amplify sgRNA cassettes for sequencing, and sgRNA enrichment was inferred with MAGeCK. **(F)** Scatter plots representing Robust Rank Aggregation (RRA) positive scores of two technical replicates for the first challenge round, showcasing sgRNA enrichment at gene-level (plotted as −log_10_(pos|score). GPI-anchored proteins are colour-coded (red, light red, dark blue) and putative GPI-anchor biosynthesis enzymes among the top 50 hits are labelled in blue. **(G)** Schematic visualisation of the three *Ae. albopictus* GPI-anchored membrane proteins identified in the CRISPR screen; Lachesin, Lachesin-like, and Fasciclin-1. Domains were predicted with InterProScan. In the inset, the canonical GPI-anchor structure (51) indicates the sites and chemical moieties involved in reactions catalysed by the top PIG enzymes identified in the screen. *P = < 0.05, **P = < 0.01.

First, we confirmed that both RprKR-expressing CHIKVs cause apoptosis in C6/36-derived mosquito cells upon infection. We assessed caspase-3/7 activation over time by live cell imaging using a DEVD-coupled nuclear dye that is sensitive to caspase-3/7 cleavage. As expected, wild-type (WT) CHIKV did not induce apoptotic cell death within the monitored time frame (**Fig 1B and 1C**). In contrast, mosquito cell cultures inoculated with RprKR-expressing CHIKVs showed enhanced caspase activation from 12 hpi onwards in combination with visible cell death (**Fig 1B and 1C**). Second, we assessed viral replication fitness of both RprKR viruses as compared to WT CHIKV. In mammalian BHK-21 cells, CHIKV 2SG RprKR reached infectious virus titers comparable with those of WT CHIKV, while CHIKV RprKRorf reached approximately 100-fold lower titers (**Fig 1D**). In C6/36 mosquito cells, both RprKR-expressing CHIKVs displayed reduced replication fitness compared with WT CHIKV and, again, CHIKV RprKRorf showed greater attenuation (**Fig 1D**). Based on these results, we selected the least attenuated CHIKV-RprKR virus, i.e. CHIKV 2SG-RprKR, for the subsequent genome scale loss of function CRISPR screen.

### CRISPR screen identifies GPI-anchored proteins and GPI-biosynthesis enzymes

Insect cells are largely refractory to lentiviral transduction, which is a delivery method routinely used in mammalian cell CRISPR screening. To enable genome-scale CRISPR screening in *Ae. albopictus* cells, we used our previously established strategy (30) to generate a “CRISPR-ready” C6/36HE8C cell line expressing Cas9 and harbouring a ΦC31 docking site for recombination-mediated integration of sgRNA cassettes. We designed a membrane protein-focused sgRNA library, based on the *Ae. albopictus* Foshan FPA (AalbFP1.0) genome annotation (41). The library included all predicted transmembrane and GPI-anchor containing proteins, containing 47,677 sgRNAs targeting 6,361 protein-coding genes, with each gene represented by up to 10 sgRNAs (**Fig S1A, S1B, and Supplementary Data 1**). The library was cloned into the attB library donor vector and delivered to the CRISPR-ready C6/36HE8C cell line by ΦC31 recombinase-mediated cassette exchange (RMCE) (**Fig 1E**). Following four weeks of puromycin selection of stable integrants, the knockout cell pool was inoculated with CHIKV 2SG-RprKR at MOI:0.001. Surviving cells were expanded and re-challenged at increasing MOIs during subsequent screening rounds. After each round, genomic DNA was isolated and assessed for gene-level sgRNA enrichment relative to uninfected controls.

In the first round of our screen, we identified Lachesin (VectorBase: AALFPA_064030) as one of the top hits (**Fig 1F and Supplementary Data 2**). Lachesin is a GPI-anchored cell adhesion protein that has been associated with neurogenesis and tracheal system development in insects (42-44) (**Fig 1G**). In contrast to our expectations, enrichment scores decreased in the subsequent challenge rounds, but *lachesin* remained highly enriched (**Fig S1C and S1D**). Other GPI-anchored proteins that we identified, albeit with lower enrichment (Robust Rank Aggregation (RRA)) scores, were Lachesin like and Fasciclin-1 (**Fig 1F and 1G**). Lachesin-like has 25.95% amino acid identity with Lachesin and is not yet characterised. Fasciclin-1 is a cell-adhesion molecule implicated in insect neuronal development (45-47). Lachesin and Lachesin-like are both IgSF proteins containing three Ig-like domains (**Fig 1G**). According to sequence patterns and length, three overarching types of Ig-like domains exist; constant (C-set), variable (V-set), and intermediate (I-set) (48). Lachesin domain 1 (D1) is predicted to be V-set, D2 C-set, and D3 I-set. Lachesin-like D1 is predicted to be I-set and D2 and D3 are likely constant. Fasciclin-1 contains four FAS1 domains, which is an ancient cell-adhesion domain with multiple binding interfaces. In adult *Ae. aegypti* mosquitoes, *lachesin* transcript is broadly expressed, as based on single-nuclei transcriptomics performed by Goldman *et al*. (49) (Fig S2). This includes cells of the midgut and salivary glands, where CHIKV replicates. In that same dataset, *lachesin-like* transcript is less abundant, whereas *fasciclin-1* is ubiquitously expressed with highest abundance in neuronal tissues. Apart from these three GPI-anchored proteins, multiple enzymes involved in various steps of GPI-anchor biosynthesis were among the top hits (**Fig 1F and 1G**). Most prominently, sgRNAs targeting phosphatidylinositol N-acetylglucosaminyltransferase subunit A (*PIG-A*, VectorBase: AALFPA_075259) and GPI mannosyl transferase 3 (*PIG-B*, VectorBase: AALFPA_068932) were enriched.

### Lachesin mediates entry of CHIKV and other arthritogenic alphaviruses

As the GPI-anchored proteins identified in our screen are adhesion molecules predicted to be expressed on the plasma membrane, we hypothesised that they might be involved in CHIKV entry. To test this hypothesis, we utilised the human lymphoblast cell line K562, which does not express bona fide alphavirus entry receptors, but supports all post-entry steps of the CHIKV replication cycle. Refractory K562 cells can become susceptible to alphavirus infection upon receptor complementation. Here, we transduced K562 cells to express positive control human MXRA8 or FLAG-tagged *Ae. albopictus* Lachesin, Lachesin-like, or Fasciclin-1. Surface expression of each construct was confirmed by antibody staining and flow cytometry (**Fig 2A**). Next, we inoculated the cells with CHIKV of the ECSA and West African (WA) lineages carrying a GFP reporter under a subgenomic promoter and followed GFP expression over time by live cell imaging (**Fig S3A**). The two CHIKV strains infected K562 that ectopically express MXRA8, whereas no infection was observed in cells transduced with an empty lentiviral vector (EV), as expected (**Fig 2B and 2C**). The CHIKVs also efficiently infected Lachesin-expressing cells. In contrast, Lachesin-like and Fasciclin-1 expression did not render K562 susceptible to CHIKV infection. These data suggest that out of the three GPI-anchored hits, only Lachesin functions as CHIKV entry factor, therefore representing a strong candidate entry receptor.

**Figure 2.**
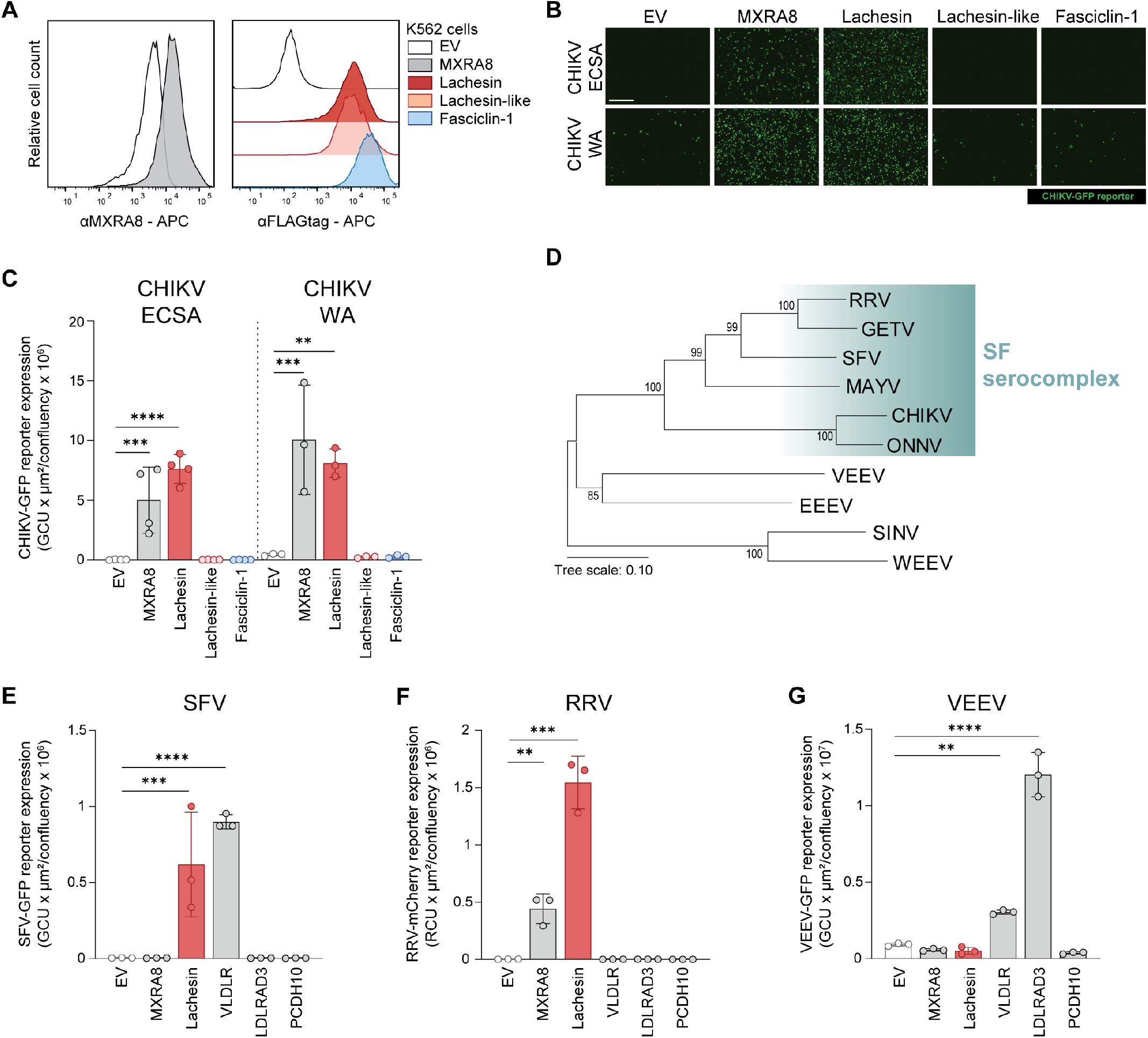
Lachesin mediates entry of CHIKV and other arthritogenic alphaviruses. **(A)** Surface expression of human MXRA8, or FLAG-tagged *Ae. albopictus* Lachesin, Lachesin-like, and Fasciclin-1 in transduced human K562 cells, as assessed with flow cytometry. As negative control, K562 cells transduced with an empty lentiviral vector (EV) were included. APC = Allophycocyanin. **(B)** CHIKV reporter virus infection (East-Central-South African (ECSA) and West African (WA) strains) at MOI:1 in K562 cells expressing human MXRA8, or *Ae. albopictus* Lachesin, Lachesin-like, or Fasciclin-1. GFP expression, encoded under a subgenomic promoter in the viral genome, was assessed by live cell imaging. Representative images are from 26 hpi. Scale bar = 400 µm. **(C)** Quantification of CHIKV reporter virus GFP expression at 26 hpi, expressed as integrated intensity (GCU x µm2/confluency), of the experiment described in **(B). (D)** Phylogenetic tree of full-length E1 protein sequences from selected alphaviruses. Tree branch lengths are proportional to the number of amino acid substitutions per site. Bootstrap support values are indicated at the nodes. GETV = Getah virus. **(E-G)** Infection with SFV4 **(E)**, RRV T48 **(F)**, and VEEV TC-83 **(G)** reporter viruses at MOI:1 in K562 cells expressing human MXRA8, VLDLR, LDLRAD3, PCDH10, *Ae. albopictus* Lachesin, or transduced with empty vector. Reporter expression was assessed by live cell imaging at 26 hpi and expressed as integrated intensity (GCU or RCU x µm2/confluency). Data in (C, E-G) are mean ± SD of three to four biological replicates. One-way ANOVA with Dunnett’s multiple comparisons test. **P = < 0.01, ***P = 0.001, ****P < 0.0001.

To test whether other alphaviruses can also use Lachesin to enter cells, we inoculated K562 cells with the arthritogenic alphaviruses SFV and RRV, which belong to the SF serocomplex, and with the encephalitic alphavirus VEEV (**Fig 2D**). As positive controls, we included K562 cells ectopically expressing the respective viruses’ entry receptors, i.e. VLDLR, MXRA8, and LDLRAD3. As additional negative control, PCDH10-expressing cells were included. Apart from infecting cells expressing the bona fide human receptors, i.e. VLDLR for SFV and MXRA8 for RRV, both SFV and RRV could use mosquito Lachesin to enter the K562 cells (**Fig 2E,F** and **S3B**). Contrastingly, VEEV did not infect Lachesin-expressing cells, whereas LDLRAD3 and to a lesser extent VLDLR expression rendered K562 cells susceptible (**Fig 2G and S3B**). Interestingly, previous reports did not observe VLDLR-mediated VEEV entry into K562 cells, but it is known that VEEV can bind the LA3 repeat of VLDLR (51). Altogether, we show that multiple arthritogenic alphaviruses engage Lachesin to enter cells.

### Lachesin is essential for CHIKV infection of mosquito cells

Next, we sought to confirm our findings in mosquito cells. We used *Ae. albopictus* C6/36HE8C cells in our CRISPR screen. However, C6/36 cells do not have functional RNA interference (RNAi) machinery (52, 53), which renders them inadequate for conventional dsRNA-mediated gene silencing approaches. Therefore, we silenced our GPI-anchored hits in the RNAi-competent *Ae. aegypti* Aag2-AF5 and *Ae. albopictus* U4.4 cells. As non-targeting control, cells were transfected with dsRNA targeting the bacterial *lacZ* gene, encoding β-Galactosidase. As positive controls, we included dsRNA targeting *argonaute 2* (*ago2*), the main component of the antiviral RNAi pathway, and dsRNA targeting the GFP reporter that is present in the virus. First, we confirmed efficient mRNA knockdown in dsRNA-transfected mosquito cells by RT-qPCR, using the housekeeping reference gene *ribosomal protein S17* (*RPS17*) for normalisation. Transcript abundance of *lachesin-like* was very low in both cell lines (**Fig 3A**), which corresponds to the limited read counts in the *Ae. aegypti* cell atlas (**Fig S2**). Transcript silencing was successful for all targets in both cell lines, except for *lachesin-like* in U4.4 cells, where transcript levels approached the lower limit of detection for reliable RT-qPCR quantification (**Fig 3A**).

We inoculated the dsRNA-transfected mosquito cells with CHIKV-GFP (ECSA-lineage) and assessed GFP-based infection levels by flow cytometry. As expected, silencing of the antiviral *ago2* in both mosquito cell lines increased infection significantly, whereas transfection of *GFP* targeting dsRNA effectively reduced the number of GFP-positive cells (**Fig 3B**). Knockdown was less efficient in U4.4 than in Aag2-AF5 cells, as *GFP* dsRNA reduced the number of GFP-positive cells to 3.7 ± 1.2% (mean ± SD) of control in Aag2-AF5 cells and to 29 ± 11.5% of control in U4.4 cells. Yet, silencing of *lachesin* resulted in a significant reduction in CHIKV infection of both Aag2-AF5 and U4.4, to 10.3 ± 5.1% and 36.7 ± 9.1% of control cells, respectively (**Fig 3B**). Silencing of *lachesin-like* or *fasciclin-1* did not affect CHIKV infection of *Aedes* spp. cells.

**Figure 3.**
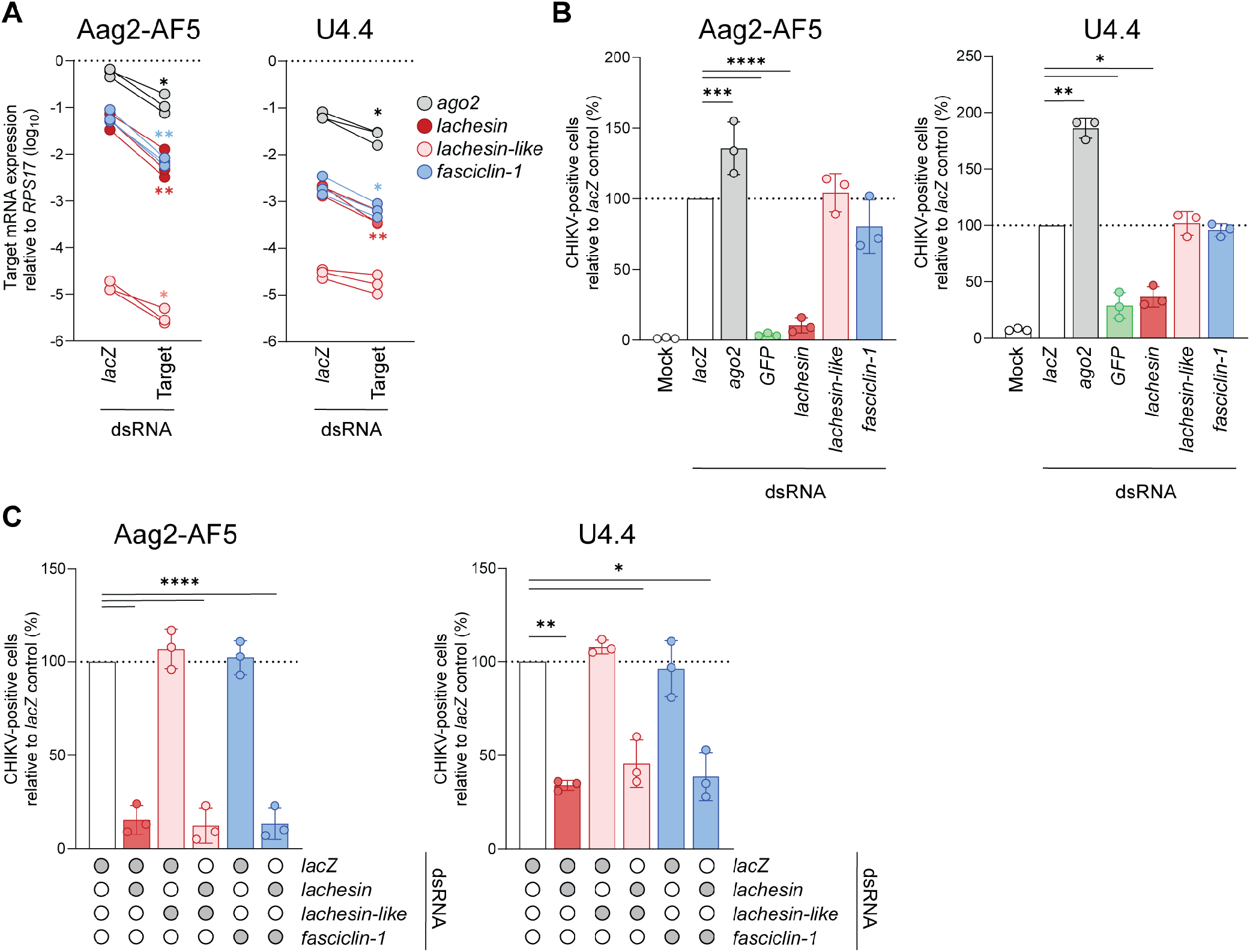
Lachesin is essential for CHIKV infection of mosquito cells. **(A)** Residual target gene expression upon dsRNA-mediated knockdown in *Ae. aegypti* Aag2-AF5 cells or *Ae. albopictus* U4.4 cells. Cells transfected with non-targeting dsRNA (negative control *lacZ*) were compared to cells transfected with dsRNA targeting the indicated genes. Gene expression was determined by RT-qPCR and normalised to the reference *ribosomal protein S17* (*RPS17*). Data are means of technical duplicates from each shown biological replicate. Paired t-tests. **(B)** Aag2-AF5 or U4.4 cells were transfected with dsRNA targeting the indicated genes or targeting *GFP* encoded in the CHIKV reporter virus. Cells were then inoculated at MOI:1 with CHIKV-GFP (ECSA-lineage) and the GFP-positive cell population was assessed at 24 hpi using flow cytometry. Infection levels were normalised to the non-targeting *lacZ* control. **(C)** Aag2-AF5 or U4.4 cells were transfected with a 1:1 mixture of dsRNA targeting the two indicated genes. Next, cells were inoculated with CHIKV and analysed as in **(B).** Data in **(B, C)** are mean ± SD of three biological replicates. RM-ANOVA with Dunnett’s multiple comparisons test. *P = < 0.05, **P = < 0.01, ***P = 0.001, ****P < 0.0001.

Since Lachesin appears to be the primary mediator of mosquito cell infection, the contribution of additional factors may be obscured in a single-target knockdown setting. Therefore, we next silenced *lachesin* and *lachesin-like* or *fasciclin-1* simultaneously. Target knockdown was significant for each co-transfected condition, except for the low-abundance *lachesin-like* (**Fig S4**). Simultaneous silencing of *lachesin* with neither of the other GPI-anchored proteins reduced CHIKV infection more than *lachesin* silencing alone (**Fig 3C**). These results emphasise that, while Lachesin-like and Fasciclin-1 appear dispensable, Lachesin is essential for CHIKV infection of *Aedes* spp. mosquito cells.

## Discussion

Arboviruses are responsible for recurrent outbreaks affecting millions of people worldwide. As alphaviruses alternate between vertebrate and invertebrate hosts and rely on mosquitoes for transmission, understanding which mosquito host factors enable viral infection is essential for developing new strategies to limit their spread. Although several bona fide receptors have been identified in mammalian hosts (24, 26-28), their mosquito counterparts remain largely unknown. Here, we coupled a genome-scale CRISPR knockout screen in *Aedes* cells with a pro-apoptotic positive selection strategy, identifying the GPI-anchored IgSF protein Lachesin as a critical host factor for CHIKV infection. Ectopic expression in refractory mammalian cells and RNAi-mediated silencing across two mosquito species demonstrated that Lachesin functions as a candidate entry receptor for CHIKV, SFV, and RRV. In parallel, Plung *et al*. used the same mosquito cell CRISPR screening platform to perform a single-round infectious CHIKV-reporter virus particle (RVP) CRISPR screen, leading to the independent identification and validation of Lachesin as mosquito receptor for arthritogenic alphaviruses (54). Together, these findings provide robust evidence for Lachesin as a mosquito entry factor for CHIKV and establish a versatile framework for dissecting vector-alphavirus interactions.

### Strengths and limitations of the screening strategy

Unlike mammalian cells, mosquito cells typically survive arboviral infections and frequently establish persistent and non-cytolytic infections (33-36, 55), rendering conventional survival-based positive selection CRISPR screens ineffective. We overcame this limitation by engineering CHIKV to express the *Drosophila* pro-apoptotic protein Reaper, converting viral infection into a selectable phenotype in which only cells carrying protective genetic perturbations survive. This strategy enabled the application of stringent positive selection CRISPR screening to mosquito cells and provides a simple, cost-effective, and broadly applicable screening platform, particularly in high-containment settings where conventional cell sorting approaches can be challenging to implement.

Inoculation of mosquito cells with CHIKV 2SG-RprKR at MOI:0.001 resulted in cell death of virtually all cells within two days, creating an extreme genetic bottleneck. The enrichment of Lachesin, associated GPI-anchoring enzymes, and hits such as the cholesterol transporter Niemann-Pick disease type C1 (NPC1, VectorBase: AALFPA_058960) suggests that the selection strategy preferentially identified host factors acting at early stages of infection. We propose that, even when later stages of the viral life cycle are impaired by disruption of proviral replication factors, residual viral RNA replication is sufficient to drive subgenomic transcription and expression of the Reaper transgene to induce apoptosis. Additionally, viral entry mechanisms that rely on redundancy among multiple host factors may be challenging to capture with this type of selection-based screen. In such cases, depletion of individual components results in a partial reduction in infection rather than complete restriction, allowing these factors to escape detection under a highly stringent pro-apoptotic selection strategy. Accordingly, the identification of Lachesin as a top-enriched hit suggests that this factor is critical and non-redundant for CHIKV infection.

Surviving cell populations were expanded for subsequent re-challenges with CHIKV 2SG-RprKR with the goal of increasing the signal-to-noise ratio. However, we observed a stark reduction in sgRNA enrichment over time (**Fig S1C and S1D**). Moreover, no cell death was observed in subsequent challenge rounds. The latter could reflect the sole presence of cells containing a knockout of a CHIKV dependency factor, but this does not match a decrease in enrichment scores. Therefore, it is more likely that CHIKV 2SG-RprKR may have progressively lost expression of the Reaper cassette, as has been reported previously for SINV-Reaper *in vivo* (38). Accumulation of these non-apoptotic viral variants could reduce selection stringency and increase background noise. For future applications, optimisation of the challenge virus is warranted to achieve optimal balance between replicative fitness and genomic stability. In addition, initiating the screen at a higher MOI than 0.001 may reduce reliance on secondary rounds of infection and thereby minimise the selective advantage of revertant viruses. Notably, despite the decline in enrichment observed during subsequent challenge rounds, the initial selection led to an exceptionally strong enrichment of the top candidate genes. The highest-ranking hits, including *PIG-B* (RRA positive score = 1.06 × 10^-15^) and *lachesin* (1.14 × 10^-12^), achieved RRA positive scores four to seven orders of magnitude lower than those reported using the complementary single-round CHIKV-RVP screening strategy (54), where the strongest hit, *lachesin*, reached a maximum RRA positive score of 1.95 × 10^-8^. These findings demonstrate that the pro-apoptotic selection strategy provides exceptionally stringent and sensitive enrichment during the initial screening round, making it particularly well suited for robust prioritization of candidate host factors.

### Lachesin as a CHIKV entry factor in mosquitoes

Lachesin is GPI-anchored, which sets it apart from transmembrane helix-containing alphavirus entry receptors identified in mammals, i.e. MXRA8, VLDLR, ApoER2, LDLRAD3, and PCDH10 (24, 26-28). Interestingly, when the cytoplasmic and transmembrane domains of MXRA8 were replaced by a GPI-anchor, it could still mediate CHIKV uptake (56). Previously, other GPI-anchored virus entry factors have been identified. GPI-anchored folate receptor-α (FOLR1) was initially proposed as a filovirus receptor (57). However, subsequent studies established multipass transmembrane protein NPC1 as the essential intracellular receptor for Ebola and Marburg viruses (58, 59), while FOLR1 is no longer considered physiologically relevant. The GPI-anchored complement regulatory factor Decay-Accelerating-Factor (DAF/CD55) is an attachment factor for Enterovirus 70 (60, 61), several echoviruses (62), and coxsackieviruses B3 and A21 (63, 64). Following binding of coxsackievirus to DAF, the virion relocates to the tight junctions where it interacts with its bona fide receptor coxsackievirus and adenovirus receptor (CAR) to trigger viral uncoating (65). Interestingly, virion binding to DAF on the apical face of polarised cells leads to kinase signalling, which triggers lateral translocation of virions to the CAR-expressing tight junctions. Thus, despite the absence of a cytoplasmic signalling domain, binding of GPI-anchored proteins by extracellular ligands can lead to intracellular signalling events, which are often cluster-driven (66). Whether Lachesin is a bona fide mosquito entry receptor for CHIKV or whether it is an entry co-factor like DAF, will be subject of further studies. However, the fact that K562 cells became susceptible to CHIKV upon Lachesin overexpression strongly suggests that Lachesin functions as an entry receptor. Furthermore, Plung *et al*. demonstrated that Lachesin physically interacts with CHIKV glycoprotein, a key characteristic of viral entry receptors (54).

Lachesin belongs to the IgSF, which comprises many virus receptors (67), including MXRA8 (24). While Lachesin contains three Ig-like domains, the MXRA8 ectodomain contains two strand swapped Ig-like domains oriented in a disulfide-linked head-to-head arrangement (25). MXRA8 binds a conformationally defined surface on CHIKV, inserting into a cleft formed by two adjacent E2–E1 heterodimers and extending its interactions to a neighbouring trimeric spike (25). Lachesin is predicted to adopt an Ig-like fold resembling MXRA8 (**Fig S5A-C**), but structural studies, such as cryo-electron microscopy, will be required to resolve its topology and the interface between Lachesin and the CHIKV glycoproteins.

The biological function of Lachesin has been explored in *D. melanogaster*, where it is important for neuronal and tracheal development (43, 44), but its function in mosquitoes remains enigmatic. *Drosophila* Lachesin has 72.58% amino acid identity with *Ae. albopictus* Lachesin and is a homophilic adhesion molecule (43, 44). In *Drosophila*, Lachesin is part of the Septate Junctions (SJs), which are functional equivalents of vertebrate tight junctions and are found in epithelia and glia that form the blood–brain barrier (43, 68). Using immuno-electron microscopy, Lachesin was found to localise to the SJs of the developing tracheal tubes, basal to the apical rim of the Zonula Adherens (43). Our Alphafold3 modelling of *Ae. albopictus* Lachesin generated a low-confidence dimer model in which the D1 domains of two Lachesin molecules are positioned in a head-to-head arrangement (**Fig S5D**). Although this observation should be interpreted cautiously, it is consistent with a possible role for Lachesin in homophilic cell–cell adhesion. In mammalian cells, several virus receptors and entry factors are components of cell junctions. These include, among others, the above-mentioned CAR, the junction adhesion molecule-A as a receptor for reovirus (69, 70), and the tight junction molecules claudin-1 and occludin, which are entry factors for hepatitis C virus (HCV) (71, 72). While virus-CAR interactions can disrupt intercellular adhesion and barrier function (73), reovirus and HCV engagement of junction proteins does not impair barrier function (74). The mode of engagement of Lachesin by CHIKV in intact mosquito tissues is an important aspect of future work. A strong impairment of epithelial barrier functions in mosquitoes by Lachesin engagement is, however, unexpected as CHIKV can persistently infect mosquitoes without obvious pathology. Surprisingly, Lachesin has been reported as Dengue virus entry factor in mosquitoes (75), but the protein analysed in that study does not correspond to current *lachesin* annotations.

Upon infection of mammalian cells, CHIKV attaches to cells via PS-receptors and glycosaminoglycans, eventually binding an entry receptor that leads to internalisation and pH-dependent fusion (17, 18, 76, 77). The consensus is that clathrin-mediated endocytosis (CME) is the primary mode of uptake, followed by pH-dependent fusion of the viral envelope with the membrane of early endosomes (56, 76, 78, 79). However, clathrin-independent mechanisms have also been reported, and exact mechanisms appear cell-type dependent (80). In mosquito C6/36 and Aag2 cells, attachment is independent of PS-receptors and glycosaminoglycans (77). Similarly to mammals, membrane fusion in mosquito cells is triggered by endosomal acidification, although it occurs in late rather than early endosomes (81-83). CHIKV entry into C6/36 cells is receptor- and cholesterol-dependent and CME and Eps15 are likely involved (82, 83). Previous studies attempted to identify CHIKV receptors in mosquitoes by combining virus overlay protein binding assays (VOPBA) with mass spectrometry (84-86). However, validation of these candidates remained limited, and some hits were subsequently shown to result from experimental artefacts (77). With the knowledge that Lachesin is involved in CHIKV-entry into *Aedes* spp. cells, detailed studies are warranted to elucidate the exact mechanisms of virus attachment, endocytosis, and fusion.

In addition to Lachesin, the cell-adhesion molecules Lachesin-like and Fasciclin-1 were enriched in the CRISPR screen, but we observed no CHIKV-related phenotype during follow-up experiments. Phenotype validation was performed in a K562-overexpression system and in Aag2-AF5 and U4.4 cells. As these differ from CRISPR-ready C6/36HE8C cells, the alternative hits may represent CHIKV host dependencies specific to this clone. An additional consideration is the possibility that multiple donor plasmids entered the same cell during RMCE, resulting in simultaneous disruption of more than one gene. Under these circumstances, the phenotype associated with an enriched sgRNA may, in some cases, reflect the contribution of an additional, unrecognised mutation. Nevertheless, the independent enrichment of multiple sgRNAs targeting the same genes, including *lachesin* (8/10 sgRNAs), *lachesin-like* (2/3 sgRNAs), and *fasciclin-1* (6/10 sgRNAs), suggests that these loci are reproducibly associated with the selected phenotype. Future implementations of the platform could further reduce this source of noise by incorporating unique molecular identifiers (UMIs) to improve genotype– phenotype linkage. Moreover, by adopting integration strategies, such as intAC (87), which transiently suppress Cas9 activity during library integration, only stably integrated sgRNA cassettes would drive genome editing. The strong enrichment of *lachesin* and its validation as mosquito entry factor for CHIKV in orthogonal assays shows that the current platform already has high potential for future host factor discovery studies.

### Final remarks

In summary, we, in parallel with Plung *et al*. (54), have identified Lachesin as a candidate receptor for CHIKV in mosquitoes. Additionally, our work establishes a broadly applicable platform for positive selection CRISPR screening of non-cytolytic arboviruses in mosquito cells. This approach provides a scalable framework to systematically map mosquito host factors that govern arbovirus infection. This opens new opportunities for the development of targeted antiviral interventions and innovative vector-control strategies to mitigate the global burden of mosquito-borne diseases.

## Acknowledgements

We thank Jonathan Zirin, Lu-Ping Liu, Isabell Niedermoser, Ju Eun Yoo, Lifeng Liu, Lisa Lasswitz, and Verena Pittl for their valuable technical assistance. This article is subject to HHMI’s Open Access to Publications policy. HHMI lab heads have previously granted a nonexclusive CC BY 4.0 license to the public and a sublicensable license to HHMI in their research articles. Pursuant to those licenses, the author-accepted manuscript of this article can be made freely available under a CC BY 4.0 license immediately upon publication.

## Author contributions

Conceptualization: A.C.M.dB., E.M., N.A., N.P., G.G. Data Curation: E.M., A.C.M.dB. Formal Analysis: A.C.M.dB., E.M. Funding Acquisition: G.G., N.A.,

N.P. Investigation: A.C.M.dB., E.M., N.A. Methodology: E.M., A.C.M.dB. Resources: A.M., J.P., W.L., J.A. Software: Y.H., E.M., R.V. Supervision: G.G.,

N.P. Visualization: A.C.M.dB. Writing – Original Draft Preparation: A.C.M.dB., E.M., G.G. Writing – Review & Editing: A.C.M.dB., E.M., A.M., N.A., L.V., G.G.

## Materials and Methods

### Cell culture

The *Ae. albopictus* “CRISPR-ready” clonal cell line C6/36HE8-Ub::Cas9-2A-Neo (attP^+^ Cas9^+^; RRID:CVCL_F1ZQ; DGRC stock #363), referred throughout the text as “C6/36HE8C”, was generated as previously described (30) and maintained in Schneider’s Drosophila Medium (Gibco) supplemented with 10% (v/v) heat-inactivated foetal calf serum (FCS; Capricorn Scientific) and 1% (v/v) penicillin-streptomycin (Gibco) and 400 μg/mL Geneticin (G418; Sigma). The wild-type *Ae. albopictus* C6/36 cell line (ATCC CRL-1660) was maintained in Schneider’s Drosophila Medium (PAN Biotech) supplemented with 10% FCS, 2 mM L-glutamine (Gibco), 0.1 mM non-essential amino acids (NEAA; Gibco), 1% sodium pyruvate (Gibco), and 1% (v/v) penicillin-streptomycin. The clonal *Ae. aegypti* cell line Aag2-AF5 (ECACC 19022601) (88), kindly provided by Kevin Maringer (Pirbright Institute, United Kingdom), was maintained in GlutaMAX-supplemented Leibovitz’s L-15 Medium (Gibco) containing 10% (v/v) FCS, 10% (v/v) tryptose phosphate broth (TPB; Gibco), 0.1% NEAA, and 1% (v/v) penicillin-streptomycin. The *Ae. albopictus* U4.4 cell line (CVCL_Z820; Tick Cell Biobank, United Kingdom) was maintained in GlutaMAX-supplemented Leibovitz’s L-15 Medium supplemented with 20% (v/v) FCS, 10% (v/v) TPB, 1% (v/v) L-glutamine, and 1% (v/v) penicillin-streptomycin. All mosquito cell culture and experimental procedures were performed at 28 °C.

Baby Hamster Kidney cells (BHK-21; ATCC CCL-10) and Human Embryonic Kidney cells (HEK293T; ATCC CRL-3216) cells were maintained in high glucose Dulbecco’s Modified Essential Medium (DMEM; Gibco) supplemented with 10% (v/v) FCS, 0.1 M NEAA, 1% (v/v) L-glutamine, and 1% (v/v) penicillin-streptomycin. Human chronic myelogenous leukaemia lymphoblast K562 cells (ATCC CCL-243) were maintained in Roswell Park Memorial Institute (RPMI) 1640 Medium (Thermo Fisher Scientific) supplemented with 10% (v/v) FCS, 25 mM HEPES, and 1% (v/v) penicillin-streptomycin. Mammalian cell culture and experimental procedures were performed in a humidified incubator at 37 °C and 5% CO_2_.

### Viruses

Virus stocks of infectious cDNA (icDNA) clones from CHIKV LR2006 OPY1 GFP and CHIKV 37997 GFP (provided by G. Simmons) (89), and VEEV TC-83 GFP (provided by I. Frolov) (90) expressing GFP under the control of a subgenomic promoter (SGP) (91), and SFV4 GFP (92) and RRV T48 mCherry (provided by C. Schmidt and B. Schnierle) (93) harbouring GFP or mCherry fused to nsP3 respectively, were generated upon electroporation of plasmid DNA (SFV4) or *in vitro transcribed* mRNA (all others) in BHK-21 cells. Supernatant containing infectious virus was harvested after 2 days, clarified, and stored at −80 °C. Tissue Culture Infectious Dose 50% (TCID_50_) titers were determined by endpoint titration in BHK-21 or C6/36 cells, using reporter fluorescence at 2 dpi or cytopathic effect at 4 dpi as readout, and calculated according to Spearman-Kärber (94). Experiments with CHIKV were performed under biosafety level (BSL) 3 or 3** conditions and all other virus procedures were performed under BSL2 conditions.

Two viruses expressing *D. melanogaster* Reaper (Genbank: NP_524138.1), in which five lysine residues were replaced with arginine residues (i.e. RprKR), were constructed using the previously described icDNA backbone of CHIKV LR2006 OPY1 (95), synthetic DNA fragments (GenScript), and restriction enzyme-based cloning. First, the sequence encoding RprKR was inserted between sequences encoding human ubiquitin at the N-terminus and a tandem Myc epitope tag followed by the foot-and-mouth disease virus 2A peptide at the C-terminus. The resulting cassette was inserted in the icDNA backbone between the regions encoding the capsid protein and the E3 glycoprotein, as described previously (40), to obtain SP6-ICRES1-RprKRorf. Second, to obtain SP6-ICRES1-2SG-RprKR, the sequence encoding RprKR was inserted under the control of the natural CHIKV SGP. To ensure expression of the viral structural proteins, a duplicated copy of the SGP, corresponding to residues ™78 to +69 relative to the start site of the subgenomic RNA, was added downstream of the sequence encoding RprKR. Virus stocks were generated and titrated as described above.

### Caspase activation

To assess the induction of apoptosis upon infection, C6/36HE8-Ub::Cas9-2A-Neo cells were seeded at 20,000 cells per well in a 96-well plate, in complete medium. The next day, medium was replaced with complete medium containing the indicated CHIKV at MOI:1 and a caspase-3/7 cleavage dependent dye that allows quantification of apoptosis induction (Sartorius #4440) diluted 1:1000. Green fluorescence was monitored up to 24 hpi by the Incucyte S3 imaging platform (Sartorius) in a humidified incubator at 37 °C with 5% CO_2_. Images were captured every 2 h at 10× magnification, acquiring 4 fields of view per well. Data were analysed using the manufacturer’s Basic Analyzer tool and exported as integrated fluorescent intensity (in Green Calibrated Units (GCU) per image). At each time point, data were normalised to mock-inoculated cells to correct for photobleaching.

### Replication kinetics

BHK-21 and C6/36 cells were seeded at 500,000 cells per well in a 12-well plate, in complete medium. The next day, cells were washed once with DPBS and inoculated with CHIKV at MOI:0.001 (based on respective cell-based titers). After 2 h, cells were washed with DPBS twice and 1 mL respective complete medium was added. At indicated time points, 100 µL supernatant was harvested, frozen at −80 °C, and replenished by 100 µL fresh medium. Infectious virus titers were determined by endpoint titration in BHK-21 cells using cytopathic effect at 4 dpi as readout and expressed as TCID_50_/mL.

### Membrane-focused CRISPR knockout library

A membrane-focused CRISPR knockout library was designed to target *Ae. albopictus* genes encoding predicted transmembrane or GPI-anchored proteins. Protein-coding gene models from the *Ae. albopictus* Foshan FPA genome assembly (AalbFP1.0; VectorBase release 59) were supplemented with *de novo* predictions of membrane localisation. Transmembrane proteins were identified using DeepTMHMM v1.0.20, while GPI-anchored proteins were predicted with NetGPI 1.1. To account for potential non-canonical GPI-anchored proteins lacking an identifiable N-terminal signal peptide, the C-terminal 201 amino acids of each protein were additionally screened for predicted ω-site motifs independent of signal peptide detection. The library also included all annotated components of the GPI-anchor biosynthetic pathway, the complete mosquito kinome and phosphatome assembled from VectorBase annotations and orthology to curated *Drosophila* datasets, selected membrane-associated gene families, and a small set of positive-control genes. Inclusion of GPI-anchor biosynthetic enzymes provided a complementary functional strategy to identify GPI-anchored proteins that may escape computational prediction by disrupting surface presentation of the entire protein class. sgRNAs were designed using the CRISPR GuideXpress pipeline (https://www.flyrnai.org/tools/fly2mosquito/web/) as previously described (30). Candidate guides were prioritised according to predicted on-target efficiency, minimal off-target potential, and perfect sequence identity to the *Ae. albopictus* C6/36 genome assembly (BioProject PRJNA345486), excluding guides targeting the region that contains mismatches between the reference and the cell-line genome. Up to ten sgRNAs were selected per gene, resulting in a final library comprising 47,677 unique sgRNAs targeting 6,361 protein-coding genes together with 100 non-targeting control guides. Library oligonucleotides (109-mer; Agilent) were recovered by dial-out PCR using Q5 Hot Start High-Fidelity DNA Polymerase (NEB) and cloned by Gibson assembly into BbsI-linearised pLib6.10BN-Aaeg_774 (*GenBank accession pending; Addgene pending*) using NEBuilder HiFi DNA Assembly Master Mix (NEB). Library quality was assessed by deep sequencing of PCR-amplified sgRNA inserts (Illumina NovaSeq 6000), confirming recovery of all 47,677 designed guides. The resulting library was designated MFCRISPKO_AALB (**Supplementary Data 1**).

### CRISPR screen

To generate the knockout cell pool, C6/36HE8-Ub::Cas9-2A-Neo cells were seeded in 20x 100 mm dishes at 10,000,000 cells per dish in complete growth medium with 500 µg/mL Geneticin. The next day, the cells were at 50% confluency and transfected with a plasmid mixture containing equimolar pAeUb::ΦC31-Integrase (*GenBank accession pending; Addgene pending*) and the MFCRISPKO_AALB sgRNA donor library using Lipofectamine 3000 (Thermo Fisher Scientific). Per dish, 20 µg DNA was mixed with 40 µL P3000 reagent and complexed with 40 µL Lipofectamine 3000 in Opti-MEM. The next day, each dish was expanded into 3x 150 mm dishes in medium supplemented with 2 µg/mL puromycin. Afterwards, cells were expanded up to 8x 150 mm dishes and cultured for an additional 30 d with medium changes and reseeding at 1:10 ratio every 3 d; at each passage, total cell numbers were maintained above 1,000 cells per sgRNA to preserve library representation.

For screening, knockout-pool cells were plated at 11,000,000 cells per 150 mm dish in 4x dishes per replicate (~880-fold coverage per guide). The next day, cells were inoculated with CHIKV 2SG-RprKR at MOI:0.001 (based on C6/36 titer; designated ROUND 1). At 2 dpi, ~99% of the inoculated cells were dead as per visual inspection. Supernatant was replaced by conditioned medium every 2 d. At 7 dpi, surviving cell clusters were reseeded and expanded for re-challenge. We re-challenged with CHIKV 2SG-RprKR another three times, at MOI:0.001 on 12 d post initial challenge (dpic), at MOI:0.01 on 18 dpic (designated ROUND 2), and at MOI:1 on 25 dpic (designated ROUND 3). Following each round of CHIKV 2SG-RprKR infection, a surviving cell fraction was expanded to 2x 150 mm dishes for genomic DNA isolation. To inactivate infectious CHIKV, cell pellets were resuspended and lysed in DNA/RNA Shield (Zymo Research) after being washed twice in DPBS. Cell lysates were stored at −80 °C and genomic DNA was isolated using the Quick-DNA Midiprep Plus Kit (Zymo Research #D4075). Genomic DNA was used as the template for PCR amplification of integrated sgRNA cassettes. Amplicons were barcoded using a two-step PCR protocol (primer sequences are provided in **Supplementary Table 1**) and subjected to deep sequencing on an Illumina NovaSeq 6000. Sequencing quality metrics indicated excellent sgRNA representation throughout the screen. The plasmid library exhibited a Gini index of 0.031, which increased progressively to 0.106 after three rounds of positive selection, consistent with biological enrichment while maintaining near-complete library complexity (>99.9% of sgRNAs detected in all samples). Gene-level enrichment analysis was performed using MAGeCK (v0.5.9.5). Complete results from all screening rounds are provided in **Supplementary Data 2**.

### Generation of stable cell lines

We ordered geneblocks encoding human codon optimised *Ae. albopictus* Lachesin (VectorBase AALFPA_064030.R32171), *Ae. albopictus* Lachesin-Like (VectorBase AALFPA_041801.R1774), *Ae. albopictus* Fasciclin-1 (VectorBase AALFPA_052152.R16164; shortest transcript), and human MXRA8 (isoform 1 (Genbank NM_001282585) with C-terminal MycFLAG tag) from Integrated DNA Technologies (IDT). To allow protein detection, Lachesin and Lachesin-like were FLAG-tagged (DYKDDDK) after Ig-like domain 3, following V322 and T344 respectively; Fasciclin-1 was N-terminally FLAG-tagged after the signal peptide, between S42 and R43.

The above constructs were cloned into pWPI_IRES_Puro_Ak (from Sonja Best (Addgene #154984)) lentiviral vector. HEK 293T cells were transfected using PEI at equal mass ratio with pWPI encoding the tagged codon optimised constructs, pVSV-G encoding the vesicular stomatitis virus glycoprotein, and lentiviral packaging plasmid pCMV_ΔR8-74. Lentiviral particles were harvested as described previously (17), and added to K562 cells, followed by puromycin selection at 2 μg/mL. Alternatively, cDNAs encoding human MXRA8 (GenBank NM_032348), human PCDH10 (GenBank NM_032961), human VLDLR (GenBank NP_003374), and human LDLRAD3 (GenBank AAI43825) were cloned into lentiGuide-Puro (Addgene #52963) and used to transduce K562 cells, as described previously (26, 28, 54).

### Surface staining and flow cytometry

Cell lines were confirmed to express the transduced constructs by cell-surface immunostaining. Cells were detached in cold 0.02% EDTA in PBS and stained with 0.2 µg/mL mouse-anti-MXRA8 (MB-W040-3; MBL Life Science) or 1 µg/mL mouse-anti-FLAG (F1804; Sigma) in PBS with 1% FCS for 1 h on ice. Cells were washed twice with 1% FCS in PBS. Goat-anti-mouse-APC (A865; Invitrogen) secondary antibodies were added at 4 µg/mL for 30 min at RT. Cells were washed, and fixed with 1% PFA for 20 min on ice. Protein surface expression was determined in the live singlet gate using the BD FACS Canto II and visualised using FlowJo software v10.10.0.

### Infection of stable cell lines

K562 cells transduced with an empty lentiviral puromycin vector or transduced to overexpress *Ae. albopictus* Lachesin, Lachesin-like, or Fasciclin-1, or human MXRA8, PCDH10, VLDLR, or LDLRAD3 were seeded onto 0.01% poly-L-lysine (Sigma) coated 96-well plates (100,000 cells per well) and allowed to settle for 4 h. The medium was replaced with 100 µL infection medium (RPMI-1640 supplemented with 3% (v/v) FCS, 25 mM HEPES, 1% (v/v) penicillin-streptomycin) containing infectious reporter-alphaviruses at MOI:1 (based on BHK-21 titer). At 2 hpi, the cells were washed once with 100 µL DPBS, followed by the addition of 100 µL infection medium. Fluorescent reporter expression was assessed using the Incucyte S3 imaging platform (Sartorius) in a humidified incubator at 37 °C with 5% CO_2_. Images were captured every 4 h at 10× magnification until 26 hpi, acquiring 4 fields of view per well. Data were analysed using the manufacturer’s Basic Analyzer tool and exported as integrated fluorescent intensity (in Green or Red Calibrated Units (GCU/RCU)) per image normalised for phase confluence.

### RNA interference in mosquito cells

Unique dsRNAs (~300-400 bp) were generated against *Ae. albopictus* and *Ae. aegypti lachesin, lachesin-like*, and *fasciclin-1*; against the GFP reporter present in CHIKV-GFP (256 bp) and *Ae. albopictus* and *Ae. aegypti argonaute 2* (*ago2*, 650 bp) as positive controls for RNAi activity; and against bacterial *β-galactosidase* gene (*lacZ*, 505 bp) as non-targeting negative control. Target regions were selected within coding sequences. Amplicons were screened to exclude off-target binding (stretches of ≥ 17 bp) by BLAST in VectorBase. To create T7 *in vitro transcription* (IVT) templates, cDNA was produced from total RNA isolated from Aag2-AF5 or U4.4 cells with random hexamer primers, using the RevertAid First Strand cDNA Synthesis Kit (Thermo Fisher Scientific). Alternatively, CHIKV-GFP icDNA plasmid was used as PCR template. The template for *lacZ* T7 IVT was provided by A. Kohl (Liverpool School of Tropical Medicine). Amplicons were generated with 5′ T7 promoter-tailed primers (**TABLE S1**), gel-purified, and cloned into the pJet1.2 blunt cloning vector and Sanger sequenced. Novel gel-purified PCR amplicons were generated from the sequence-verified pJet1.2 plasmids as input for T7 IVT. We synthesised dsRNA with the MEGAscript RNAi kit (Ambion AM1626) according to the manufacturer’s instructions by co-transcribing complementary strands from opposing T7 promoters. After 4 h of dsRNA synthesis, products were denatured for 5 min at 75 °C and allowed to cool down at RT for 2 h to form duplexes. Leftover DNA template and single-stranded RNA was removed by DNAse-1 and RNAse digestion. Generated dsRNA was extracted using the MEGAscript RNAi kit column purification method and eluted in nuclease-free water.

For knockdown assays, Aag2-AF5 and U4.4 cells were seeded in complete L-15 medium at 240,000 cells per well in poly-L-lysine coated or uncoated 12-well plates, respectively. Two days later, cells were transfected with 1 µg dsRNA per well in L-15 supplemented with 2% (v/v) FCS and no antibiotics, using Cellfectin II (Thermo Fisher Scientific #10362100) at a 1:5 dsRNA:reagent (1µg dsRNA: 5µL Cellfectin II) ratio for 6 h; medium was then replaced with complete L-15 growth medium. For co-transfections, 0.5 µg of each dsRNA was mixed to reach a total of 1 µg dsRNA per transfection. Cells were assayed 72 h post-transfection for knockdown or subjected to alphavirus infection. For alphavirus infection, cultures were washed twice with infection medium (L-15 with 2% (v/v) FCS and without antibiotics), followed by CHIKV-GFP inoculation at MOI:0.5 (based on BHK-21 titer) diluted in infection medium. GFP expression in the live singlet gate was quantified by flow cytometry (BD Canto II) at 24 hpi and analysed using FlowJo software v10.10.0. Values were normalised to the non-targeting *lacZ* control within each biological replicate.

### RT-qPCR to quantify knockdown efficiency

To quantify knockdown efficiency, total RNA was extracted using the NucleoSpin RNA Mini kit for RNA purification (Macherey-Nagel), which includes an rDNAse incubation step, according to manufacturer’s instructions. Isolated RNA was diluted 1:10 in nuclease-free water and 2.5 μL of diluted RNA was used per RT-qPCR (12.5 μL total) with the iTaq Universal SYBR Green One-Step Kit (Bio-Rad). We used the following cycling program: 50 °C for 10 min; 95 °C for 1 min; 45 cycles of 95 °C for 10 s and 60 °C for 20 s. Target gene-specific and housekeeping gene *RPS17* primers were used at a final concentration of 300 nM and are listed in **TABLE S1**. Reactions were performed in technical duplicate, and amplification specificity was confirmed by melt curve analysis. Gene expression was normalised to the housekeeping gene by calculating ΔCt values (Ct_target_ − Ct_RPS17_). To visualise knockdown efficiency while preserving the normalised expression values of individual target genes, we plotted 1/(ΔCt^2^) values from non-targeting (*lacZ*) and mosquito targeting dsRNA-treated samples.

### Phylogeny

To compose the alphavirus phylogenetic tree, full length E1 protein sequences of the following viruses were used; CHIKV (AQX78118.1), MAYV (AZM66144.1), ONNV (YP_010775618.1), RRV (AAA47404.1), GETV (BAU70757.1), SFV (AKA67124.1), SINV (UDO48177.1), VEEV (YP_010775819.1), WEEV (AAF60166.1), and EEEV (ADB08675.1).

Sequences were aligned using the MUSCLE algorithm (96). Unrooted phylogenetic tree was constructed using the Neighbour-joining method (97) with Poisson correction and with node support evaluated by bootstrap analysis based on 1000 repetitions. Pairwise deletion was applied to ambiguous positions. Evolutionary analyses were conducted in MEGA12 (98).

### Computational and statistical analysis

Data were analysed using GraphPad Prism 10. Statistical methods are specified in the figure legends. Data are mean ± standard deviation (SD) of three or more biological replicates, unless indicated otherwise. Schematics were created with Biorender.com. Domain predictions were performed using InterProScan v107.0. Summary figures describing CRISPR library composition, sgRNA coverage, and screening results were generated in Python using Jupyter Notebook. Data were processed using pandas and NumPy and visualised using Matplotlib. Gene- and sgRNA-level screening results were imported directly from MAGeCK output files. Scatter plots of gene-level enrichment and sgRNA-level log_2_ fold changes were generated from MAGeCK gene_summary and sgrna_summary output tables, respectively.

---

**Figure S1.**
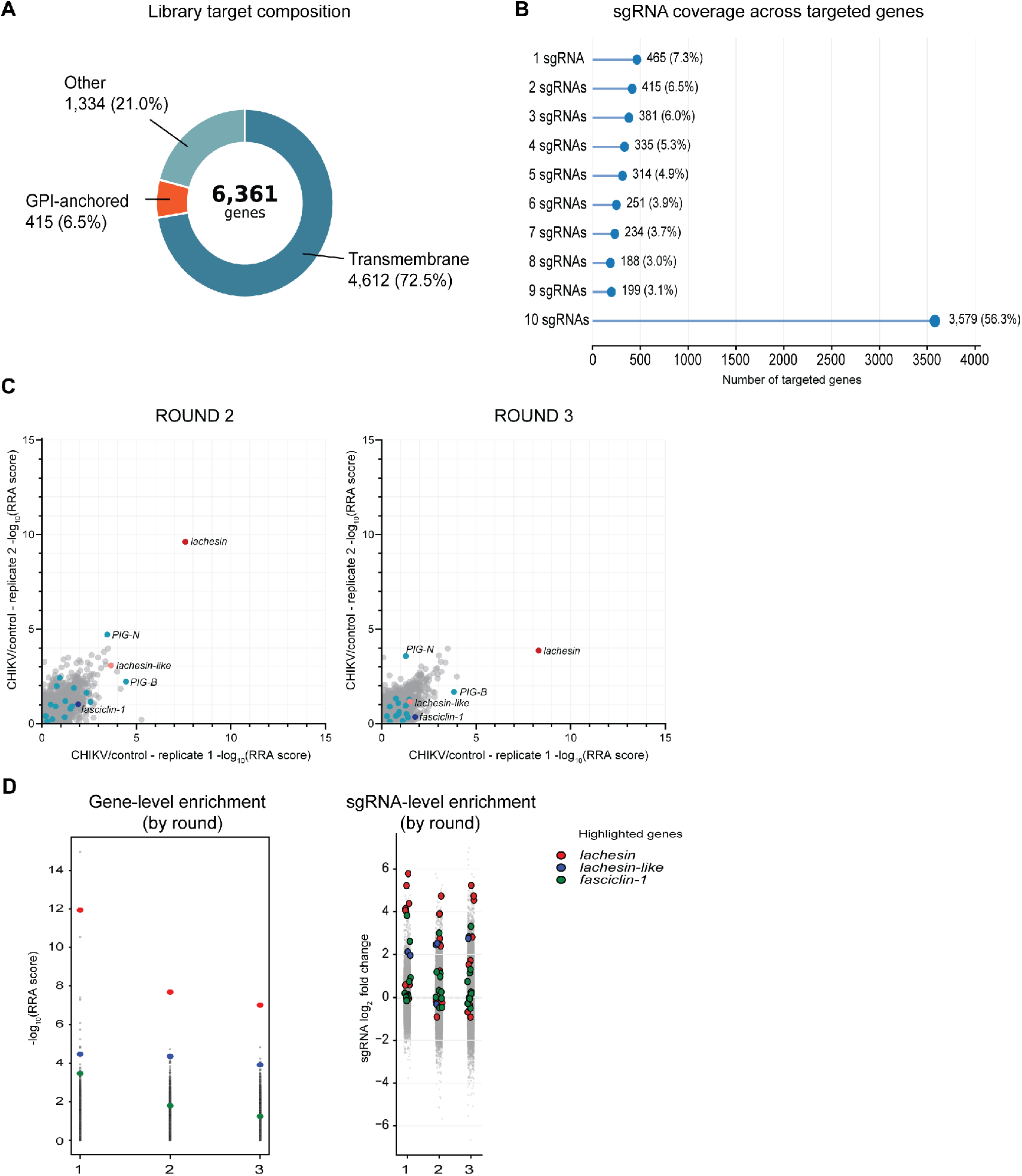
Design, composition, and screening performance of the membrane-focused CRISPR knockout library. **(A)** Composition of the membrane-focused *Aedes albopictus* CRISPR knockout library (MFCRISPKO_AALB). The library comprises 47,677 unique sgRNAs targeting 6,361 protein-coding genes, including 4,612 predicted transmembrane proteins, 415 predicted GPI-anchored proteins, and 1,334 additional genes, including kinases, phosphatases, and other non-membrane-bound proteins. **(B)** Distribution of sgRNA coverage across targeted genes. Up to ten sgRNAs were designed per gene, with 56.3% (3,579/6,361) of genes represented by a maximum of ten sgRNAs. **(C)** Reproducibility of independent biological replicates for gene-level enrichment analysis following ROUND 2 and ROUND 3 of CHIKV 2SG-RprKR selection. Each point represents one gene plotted according to its Robust Rank Aggregation (RRA) enrichment score in the two biological replicates. Selected candidate genes are highlighted. **(D)** Gene- and sgRNA-level enrichment across successive screening rounds. Left, gene-level enrichment measured as the negative logarithm of the MAGeCK RRA positive score. Right, log2 fold changes of individual sgRNAs across screening rounds. All sgRNAs are shown in grey, while sgRNAs targeting *lachesin, lachesin-like*, and *fasciclin-1* are highlighted. Multiple independently enriched sgRNAs targeting the same gene demonstrate consistent enrichment throughout the screen.

**Figure S2.**
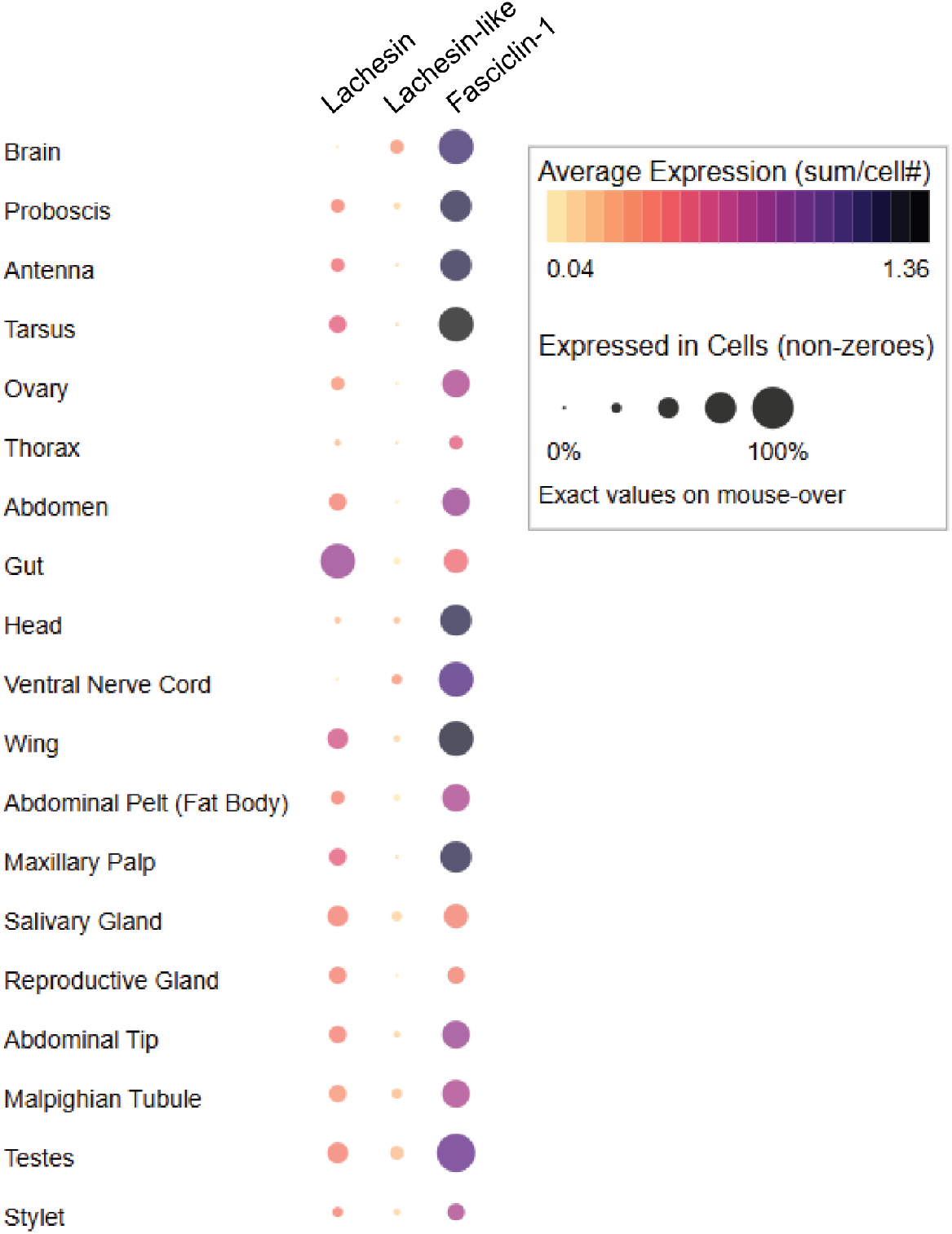
RNAseq-based mRNA expression of GPI-anchored proteins in adult *Ae. aegypti* mosquitoes. Average expression of *lachesin* (VectorBase: AAEL009295), *lachesin-like* (VectorBase: AAEL004992), and *fasciclin-1* (VectorBase: AAEL021618) across adult *Ae. aegypti* tissues, obtained from the *Ae. aegypti* single-cell atlas (49) and visualised as dot plots (UCSC Cell Browser (99)). Circle size represents the percentage of cells from each tissue that express each gene. The colours of the circles represent the average expression level of each gene across all cells from a given tissue.

**Figure S3.**
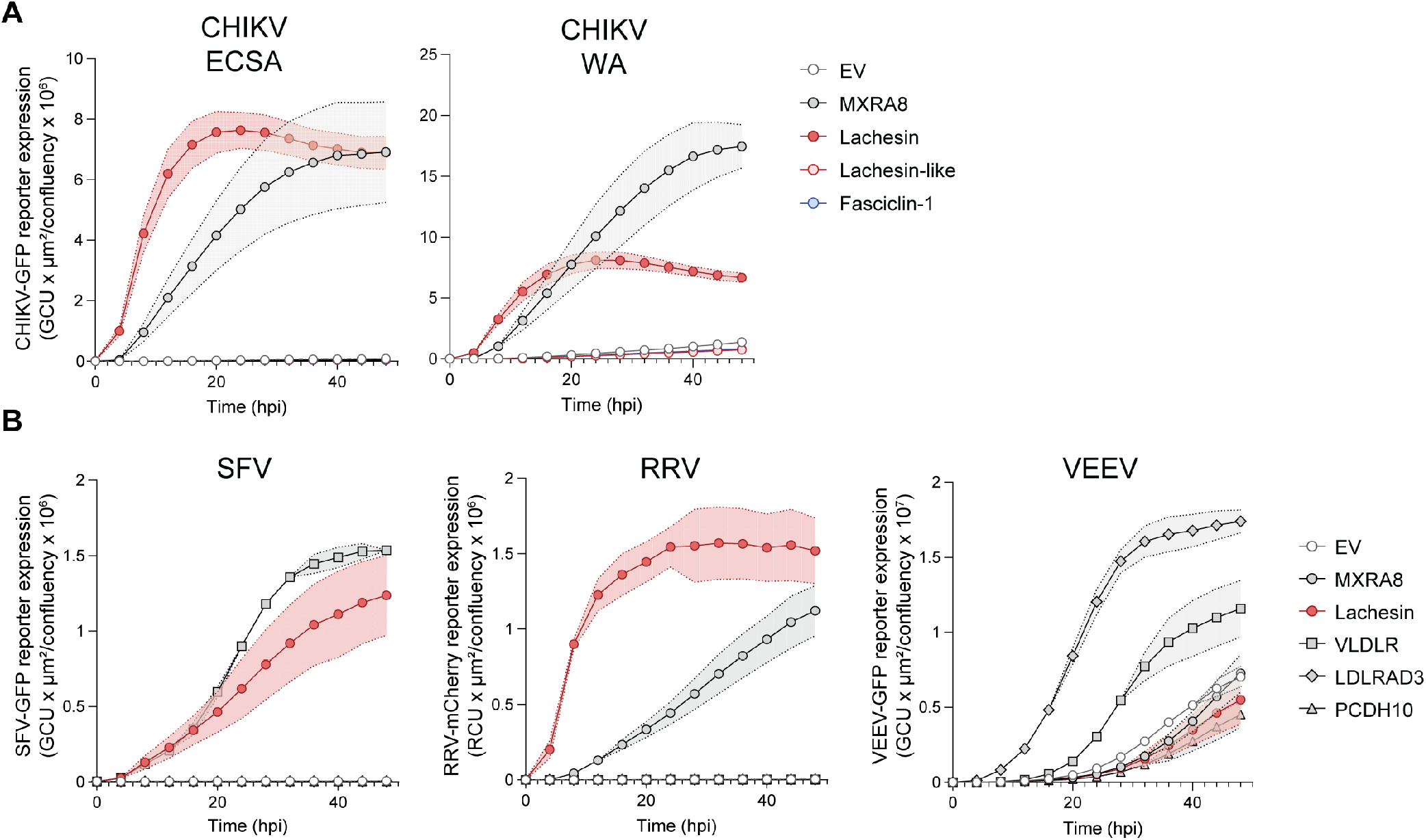
Kinetics of alphavirus infection in transduced K562 cells. **(A)** CHIKV reporter virus infection (East-Central-South African (ECSA) and West African (WA) strains) at MOI:1 in K562 cells transduced to express human MXRA8, or *Ae. albopictus* Lachesin, Lachesin-like, or Fasciclin-1. GFP expression, encoded under a subgenomic promoter, was assessed by live cell imaging and expressed as integrated intensity (GCU x µm2/confluency). EV = empty vector. **(B)** Infection with SFV4, RRV T48, and VEEV TC-83 reporter viruses at MOI:1 in K562 cells expressing human MXRA8, VLDLR, LDLRAD3, PCDH10, or *Ae. albopictus* Lachesin. Reporter expression was assessed by live cell imaging and expressed as integrated intensity (GCU or RCU x µm2/confluency). Data are mean ± SEM of two to four biological replicates.

**Figure S4.**
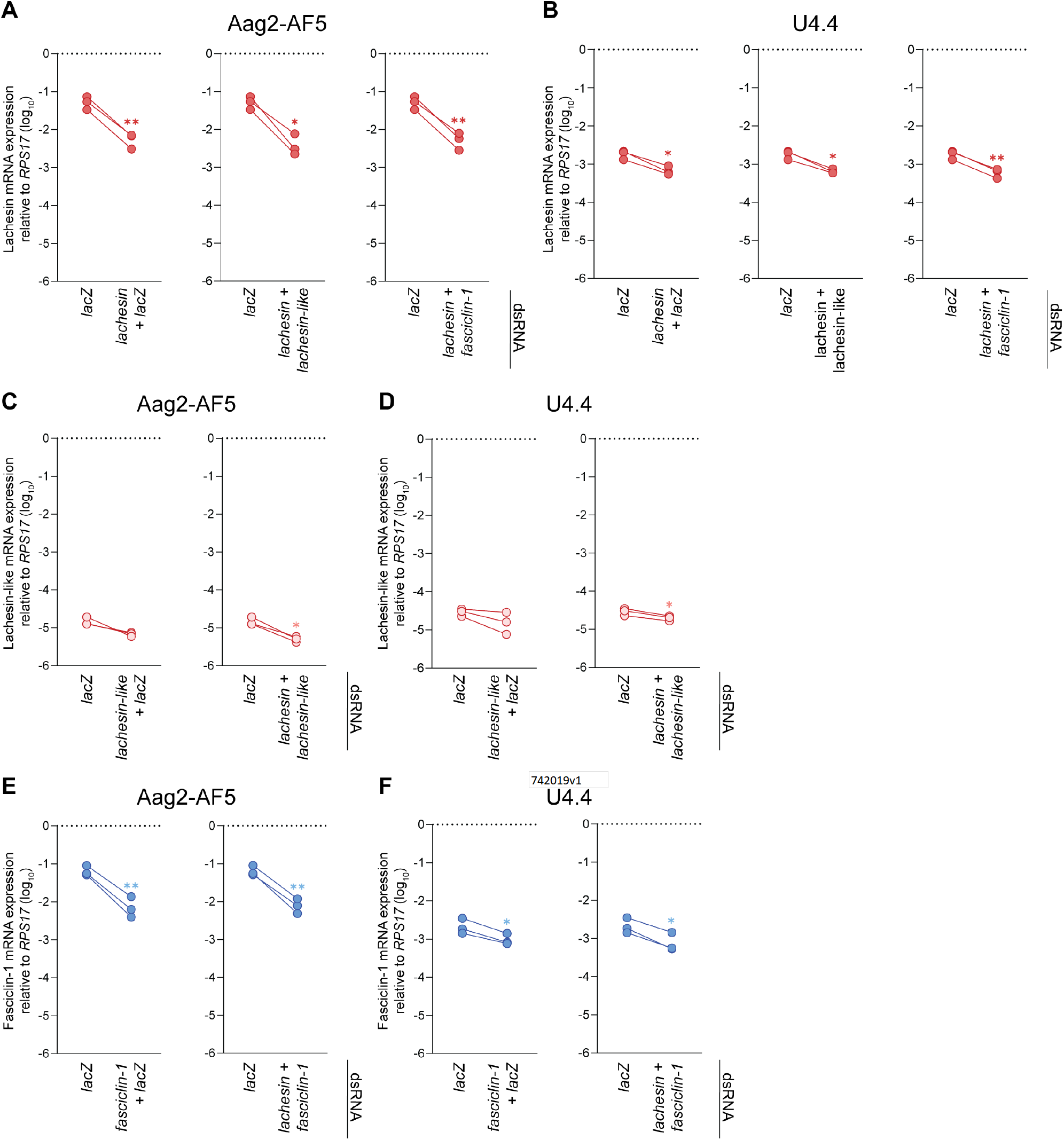
Residual expression upon simultaneous dsRNA-mediated silencing of multiple targets in mosquito cells. Residual target gene expression upon dsRNA-mediated knockdown in *Ae. aegypti* Aag2-AF5 **(A, C, E)** or *Ae. albopictus* U4.4 **(B, D, F)** cells. Cells were transfected with a 1:1 mixture of dsRNA targeting the indicated genes. Gene expression of *lachesin* **(A, B)**, *lachesin-like* **(C, D)**, and *fasciclin-1* **(E, F)** was determined by RT-qPCR and normalised to the reference *ribosomal protein S17* (*RPS17*). Data are means of technical duplicates from each shown biological replicate. Paired t-tests. *P = < 0.05, **P = < 0.01.

**Figure S5.**
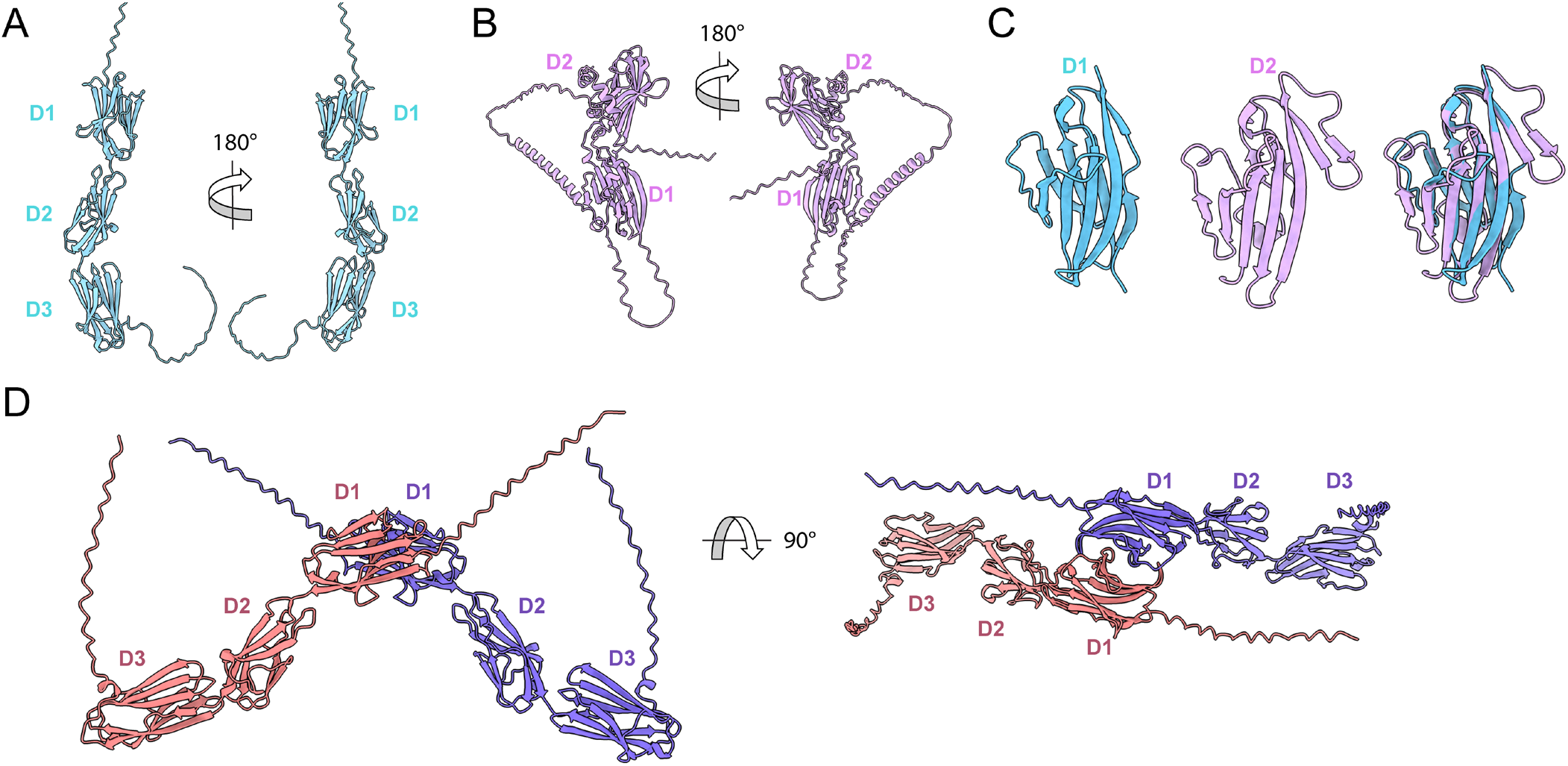
Alphafold3 modelling of Lachesin monomeric structure and potential homodimer. **(A)** 180º rotated views of Alphafold3 (AF3) model of Lachesin from *Aedes albopictus*. AF3 predicts that the protein adopts three Ig-like folds. **(B)** 180º rotated views of the AF3 prediction of full-length human MXRA8. The AF3 model closely resembles the crystal structure of mouse Mxra8 (PDB: 6JO7) (100). **(C)** Fold-based similarity alignment (ChimeraX1.12 Matchmaker) between D1 of Lachesin (Blue) and D2 of MXRA8 (Pink). **(D)** 90º rotated views of the AF3 model of the predicted D1 interaction between Lachesin protomers.

